# Standing Sentinel during Human Sleep: Continued Evaluation of Environmental Stimuli in the Absence of Consciousness

**DOI:** 10.1101/187195

**Authors:** Christine Blume, Renata del Giudice, Malgorzata Wislowska, Dominik P. J. Heib, Manuel Schabus

## Abstract

While it is a well-established finding that subject’s own names (SON) or familiar voices are salient during wakefulness, we here investigated processing of environmental stimuli during sleep including deep N3 and REM sleep. Besides the effects of sleep depth we investigated how sleep-specific EEG patterns (i.e. sleep spindles and slow oscillations [SOs]) relate to stimulus processing. Using 256-channel EEG we studied processing of auditory stimuli by means of event-related oscillatory responses (de-/ synchronisation, ERD/ERS) and potentials (ERPs) in *N* = 17 healthy sleepers. We varied stimulus salience by manipulating subjective (SON vs. unfamiliar name) and paralinguistic emotional relevance (familiar vs. unfamiliar voice, FV/UFV). Results reveal that evaluation of voice familiarity continues during all NREM sleep stages and even REM sleep suggesting a ‘sentinel processing mode’ of the human brain in the absence of wake-like consciousness. Especially UFV stimuli elicit larger responses in a 1-15 Hz range suggesting they continue being salient. Beyond this, we find that sleep spindles and the negative slope of SOs attenuate information processing. However, unlike previously suggested they do not uniformly inhibit information processing, but inhibition seems to be scaled to stimulus salience.

**Funding:** CB is supported by the Konrad-Adenauer-Stiftung e.V. and CB, MW and DPJH are supported by a grant from the Austrian Science Fund FWF (Y-777). CB, RdG, MW and DPJH are also supported by the Doctoral College “Imaging the Mind” (FWF; W1233-G17).

## 1. Introduction

Cognitive processing and task performance are well-known to vary with time of day (Dijk et al., 1992; Santhi et al., 2016; Wyatt et al., 1999). Behaviourally, these variations can readily be observed with major changes in performance paralleling the sleep-wake cycle. Beyond these within-state studies that investigated wakefulness only, we lately studied cognitive processing during the fading of consciousness, which we here define as behavioural responsiveness, that is across vigilance stages from waking to light NREM sleep (Blume et al., 2016). Specifically, during a nap we compared processing of subjectively relevant vs. irrelevant stimuli (i.e. subject’s own names [SONs] vs. unfamiliar names [UNs]) during wakefulness and non-rapid eye movement (NREM) sleep stages N1 and N2. Besides subjective relevance we additionally varied the emotional prosody of stimuli (i.e. stimuli spoken by an angry vs. a neutral voice [AV vs. NV]). Interestingly, we found evidence for preferential processing of salient stimuli (i.e. SONs and AV stimuli) not only during wakefulness, but also during light NREM sleep, with these findings suggesting not only continued processing of external stimuli, but a ‘sentinel processing mode’ of the brain during states of decreased consciousness and naturally occurring unconsciousness, that is N1 and N2 sleep, respectively. Moreover, this initial preferential processing of salient stimuli seemed to be accompanied by a subsequent inhibitory sleep-protecting process during N2 sleep that was reflected by a K-complex-like response.

In the present study we sought to replicate our previous findings on the interaction between ‘enduring brain’ or vigilance states (i.e. wakefulness, N1 and N2 sleep) and stimulus characteristics and expand them to deep N3 as well as rapid-eye-movement (REM) sleep during a full night. Beyond this, we aimed at investigating the interaction between stimulus characteristics and ‘transient brain states’, namely sleep spindles and slow oscillations representing sleep-specific electroencephalogram (EEG) phenomena in more fine-grained analyses. Sleep spindles are considered the hallmark of N2 sleep albeit they also occur during sleep stage N3. They are defined as bursts of oscillatory activity in the sigma range (11-15 Hz) with a characteristic waxing and waning shape and a duration of 0.5-3s. Slow oscillations (SOs), on the other hand, are defined as large delta waves with a first negative going wave that is followed by a positive going deflection (for criteria applied here see Riedner et al., 2007 and p. 5 of the supplementary material). Importantly, they occasionally occur during N2 sleep already, where they are often denoted K-complexes and can be considered ‘forerunners’ of or ‘sub-threshold’ SOs (Amzica & Steriade, 1997; De Gennaro et al., 2000) and are sometimes even denoted ‘peripherally evoked slow waves’ (Bellesi et al., 2014). With increasing sleep depth, the probability of occurrence of SOs strongly increases with the amount of SOs also being a criterion for deep N3 sleep.

While it is well-established that the brain is not completely shut off from the environment during sleep but continues to process external stimuli (e.g. Bastuji & García-Larrea, 1999; Blume et al., 2016; Perrin et al., 1999; Strauss et al., 2015), studies also suggest that sleep-specific oscillatory patterns, that is sleep spindles as well as SOs, can significantly alter stimulus processing. Generally, it has been suggested that during spindles the thalamus acts as a sensory filter inhibiting sensory transmission to the forebrain (Steriade, 1991). The negative or positive going slope of SOs on the other hand has been associated with changes in the probability of synaptic release at the cortical level, which could affect stimulus processing (Massimini & Amzica, 2001). In a combined EEG and functional magnetic resonance imaging (fMRI) study Schabus et al. (2012) found that responses to simple tones during NREM sleep were comparable to responses during wakefulness except for when tones were presented during a spindle or the negative going slope of a slow oscillation thereby also confirming previous findings (Dang-Vu et al., 2011; see De Gennaro & Ferrara, 2003 for an overview; Massimini et al., 2003). Likewise, in a study that looked at event-related potentials (ERPs) Elton et al. (1997) suggested that sleep spindles inhibit processing of auditory stimuli and Cote et al. (2000) additionally found the effect of sleep spindles on processing to be modulated by stimulus intensity. Specifically, they report that spindles co-occurring with more intense (i.e. louder) stimuli seemed to inhibit processing to a greater extent than was the case with less intense stimuli. Regarding slow oscillatory activity on the other hand, a pioneering study by Oswald et al. (1960) already showed that SONs evoke more K-complexes (KCs) than do unfamiliar names. Beyond this, Massimini et al. (2003) showed that evoked somatosensory EEG potentials were strongly modified not only by the presence but also by the phase of the slow oscillation. In summary, these findings strongly suggest that sleep spindles and slow oscillatory activity systematically alter stimulus processing during NREM sleep in a dynamic manner.

The aim of the present study was to investigate processing of more complex auditory stimuli (as compared to simple tones) in relation to (i) ‘enduring’ as well as (ii) ‘transient’ states of the brain. Complex stimuli were first names that varied in salience on two dimensions, namely subjective relevance (SONs vs. UNs) and familiarity or paralinguistic aspects of emotional relevance. Specifically, stimuli were uttered by a familiar voice (FV) vs. a stranger’s voice (unfamiliar voice [UFV]). Regarding the first aim, we studied stimulus processing during all ‘enduring brain states’ across the vigilance continuum (i.e. during wakefulness, N1, N2, N3 and REM sleep) irrespective of the ‘transient state’. Regarding ‘transient’ brain states, we investigated between-stimulus differences in oscillatory activity when (i) a spindle was present during stimulus presentation, when a stimulus was presented during the (ii) positive slope of a SO, (iii) during the negative slope and when (iv) stimulus presentation evoked a SO. Processing was studied by comparing oscillatory brain responses evoked by stimulus presentation in each of these cases, that is event-related synchronisation (ERS) and desynchronisation (ERD) in the delta (1-3 Hz), theta (4-7 Hz), alpha (8-12 Hz) and sigma (11-15 Hz) frequency range. Functionally, delta ERS has repeatedly been linked to attentional processes and the detection of salient or motivationally relevant stimuli (for reviews see Knyazev, 2007; Knyazev, 2012) while theta ERS has been suggested to indicate the encoding of new information as well as working and episodic memory involvement (for a review see Klimesch, 1999; Klimesch et al., 2005). Alpha ERD on the other hand is thought to reflect task demands, attentional processes and memory retrieval processes (for a review see Klimesch, 1999; Klimesch et al., 1998). Importantly, all these interpretations have been established during wakefulness and it is likely that their functional roles are different during sleep. In a previous publication, we suggested that delta and theta ERS during sleep may mirror an inhibitory sleep-protecting response following initial processing of salient stimuli as has been suggested for sigma ERS (Blume et al., 2016).

We hypothesised that oscillatory responses would mirror salience of SONs as well as FV stimuli (compared to UNs and UFV) during wakefulness. Moreover, we expected responsiveness to stimuli to vary with the ‘enduring brain state’, that is a decrease in responsiveness from wakefulness to N3 sleep. Regarding the ‘transient brain state’ we expected that when stimulus-presentation co-occurs with sleep spindles and slow oscillations the differential brain response elicited by stimulus salience would vanish. This should specifically be the case when stimulus onset coincided with the negative slope of the slow oscillation or stimulus presentation largely overlapped with a sleep spindle.

## 2. Methods and Materials

### 2.1. Participants

We recruited 20 healthy individuals for the study. Three participants were excluded from the data analysis, one dropped out after the adaptation night and two had to be excluded due to technical problems during the acquisition. The remaining sample comprised 17 participants (three males) and had a median age of 22.6 years (*SD* = 2.3 years). Prior to the study, participants gave written informed consent. Ethical consent had been obtained from the ethics committee of the University of Salzburg and the study was in accordance with the Declaration of Helsinki (World Medical Association (WMA), 1964). For more details on the study sample please see supplementary material.

### 2.2. Experimental procedure

Participants were advised to keep a regular sleep/wake rhythm with eight hours time in bed (TIB) for at least four days prior to their first visit at our sleep laboratory, which was verified with wrist actigraphy (Cambridge Neurotechnology Actiwatch ©). Participants slept in the sleep laboratory of the University of Salzburg for two nights, one adaptation night and one experimental night. For details on the experimental procedure also see Figure 1.

**Fig. 1:**
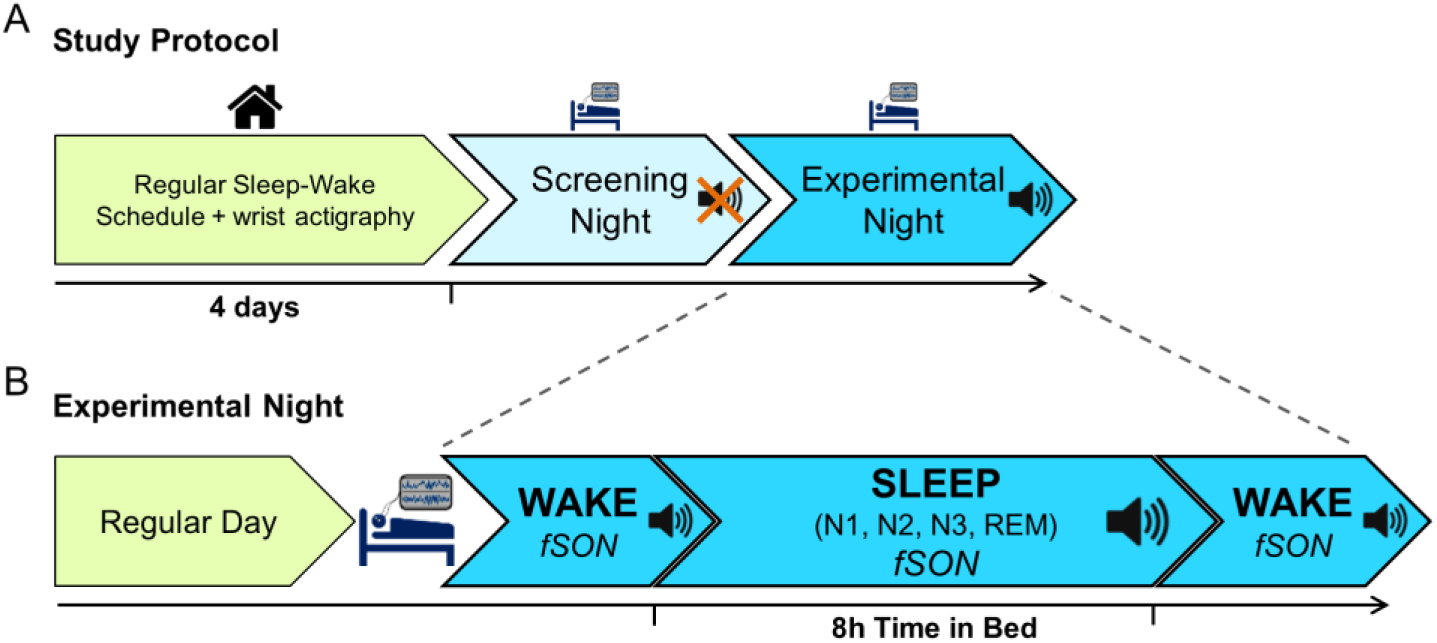
Experimental Protocol. **(A) Study Protocol.** Prior to the adaptation night in the laboratory, participants kept a regular sleep-wake schedule for four days with 8h time in bed (TIB). Adherence was verified by wrist actigraphy. During the adaptation night in the sleep laboratory polysomnography (PSG) was recorded, but no stimulation took place during sleep. **(B) Experimental Night.** The experimental night was akin to the adaptation night with wakefulness recordings preceding and following sleep. However, auditory stimulation was continued during a whole night of sleep (8h TIB).

The adaptation and experimental nights were comparable except for no auditory stimulation during sleep taking place during the adaptation night. On both nights and the following mornings participants were tested during wakefulness resulting in four wakefulness recordings per participant. The wakefulness part comprised a passive listening as well as an active counting condition, during which participants listened to the stimuli presented via in-ear headphones at a volume of approximately 65 dB. For the passive condition participants were instructed to listen attentively to the stimuli while in the active condition they were to count the number of presentations of one specific name (i.e. the target). The passive condition always preceded the active one. In this publication, we only present the results from the passive listening condition, in which participants were presented with their own name (SON) as well as two unfamiliar names (UNs) as it is the only condition that can be analysed meaningfully across ‘enduring brain’ or vigilance stages (i.e. wakefulness, NREM and REM sleep). Moreover, each name was uttered by a familiar (mother, father) and by an unfamiliar voice (lab member unknown to participant). The stimulus set was specific for each participant and all names of one stimulus set were matched regarding the number of syllables and the occurrence in the general population. During the wakefulness recording, each stimulus was presented 40 times and the interstimulus interval (ISI) was 2000ms.

Following the wakefulness recordings in the evenings, participants went to bed for an 8h±15min sleep opportunity (median sleep duration 8h 2.5min) starting at their habitual bedtime (range 8:30-11:30 pm). Participants were woken up during light NREM or REM sleep, which accounts for the jitter in the time in bed (TIB). During the experimental night, stimulation was continued and the volume was adjusted individually so stimuli were clearly audible, but participants felt they could sleep despite the stimulation. The auditory stimulation protocol was akin to the passive condition of the wake part, although during the night, the stimulus onset asynchrony (SOA) was jittered between 2.8 and 7.8 s in 500ms steps. SOA was jittered specifically in the sleep protocol as this was necessary to allow for an investigation of stimulus processing in relation to various EEG sleep phenomena (i.e. sleep spindles and slow oscillations) independent of expectation effects. SOA was not jittered during wakefulness as this would have rendered the tasks lengthy and probably too fatiguing. During the night each stimulus was presented 690 times and had the same probability of occurrence as had each SOA. For more details on the experimental procedure please see the supplementary material and Fig. 1.

### 2.3. Electrophysiological data collection and reduction

For EEG acquisition we used a 256 electrode GSN HydroCel Geodesic Sensor Net (Electrical Geodesics Inc., Eugene, Oregon, USA) and a Net Amps 400 amplifier.

#### 2.3.1. Wakefulness data

EEG data were processed using the Fieldtrip toolbox (Oostenveld et al., 2010) in Matlab (Mathworks, Natick, USA). First, the number of electrodes was reduced to 187 as the others (on the cheeks and down in the neck) contained a lot of ‘non-neural’ artefacts such as muscle artefacts and high-pass filtered at 0.5 Hz. Subsequently, eye movement artefacts were corrected using independent component analysis (ICA), data were segmented into 4s epochs (symmetrically to stimulus onset) and bad intervals were removed manually during visual data inspection. In the next step, the number of electrodes was further reduced to a final number of 173 electrodes now excluding electrodes that had initially been kept for the identification of eye and muscular artefacts. Bad channels identified during visual data inspection were interpolated and data were re-referenced to average reference. Subsequently, we randomly selected the same number of trials for each stimulus to account for imbalances in the stimulus set (only one SON, but two UNs were presented). We then applied a Morlet wavelet transformation (cycles = 3, 1-16 Hz, 1 Hz frequency steps) to each of the segments, which was followed by a baseline correction (baseline interval: −600 to 0ms relative to stimulus onset) and averaging across trials. For more details on data processing please see supplementary material.

#### 2.3.2. Sleep data

Sleep was scored semi-automatically by The Siesta Group© (Somnolyzer 24×7; cf. Anderer et al., 2005; Anderer et al., 2010; Anderer et al., 2004) according to standard criteria (American Academy of Sleep Medicine & Iber, 2007). Spindles were detected automatically during NREM sleep stages N2 and N3 at central leads using the algorithm by Anderer et al. (2005). Slow oscillations (SOs) were also detected automatically on frontal electrodes using lab-internal Matlab routines (cf. Heib et al., 2013) based on the criteria by Riedner et al. (2007) and confirmed by spot checks. For more details on the detection of spindles and SOs please see supplementary material. Pre-processing for the sleep data was essentially the same as for the wakefulness data; but we refrained from an automatic eye movement correction in order to not remove REMs. Beyond investigating processing of different stimuli across ‘enduring brain states’, that is in each sleep stage, we also investigated stimulus processing with regard to ‘transient brain states’, that is sleep spindles and SOs. To this end, we compared evoked oscillatory responses elicited by different stimuli when a spindle was present during stimulus onset (i.e. spindle offset min. 200ms after stimulus onset) or when there was a substantial overlap between a spindle and stimulus presentation (spindle onset 0-400ms after stimulus onset, i.e. spindle overlapping with at least half of the stimulus on average, cf. Suppl.Fig.1, A). Moreover, we were interested in stimulus-specific differences in the evoked slow oscillatory responses (“SO evoked”). More precisely, a SO was defined as “evoked” when the negative peak occurred between 300 and 600ms after stimulus onset (cf. Suppl.Fig.1, B1), that is the time range when the negative components of evoked K-complexes (i.e. N350 and N550) have been found to occur (Cote et al., 1999). Beyond this, we compared stimulus processing when stimulus onset was during the positive going slope of a SO (cf. Suppl.Fig.1, B2) to when stimulus onset coincided with the down-state (cf. Suppl.Fig.1, B3). For more details on data collection and analysis please refer to the supplementary material.

### 2.4. Event-Related Potentials

Although we focus on oscillatory activity in different frequency bands in the present manuscript, we provide results from event-related potential (ERP) analyses in the supplementary material (and Fig.1).

### 2.5. Statistical Analyses

Statistical analyses were performed using the cluster-based permutation approach implemented in Fieldtrip to correct for multiple comparisons that uses a Monte Carlo method for calculating significance probabilities (Maris & Oostenveld, 2007). This approach has originally been introduced by (Bullmore et al., 1999) and is referred to as the ‘cluster mass test’ in the fMRI literature (for more details please see suppl. material). Three tests were run for the main effects of *name* (SON vs. UNs), *voice* (FV vs. UFV) and the *name* × *voice* interaction with significant interaction clusters (or trends) being followed by post-hoc tests. Thus, we report three *p*-values per condition (i.e. sleep stage and interaction with sleep spindle or SO). We ran a first set of tests for the delta range that included the dimensions electrode and frequency (1-3 Hz in 1 Hz frequency steps). In the delta range, values were averaged across time (0-1000ms after stimulus onset for the WAKE condition, 0-1200ms during SLEEP) as time resolution obtained with these low frequencies was considered insufficient for an analysis in the time dimension. A second test was then run for the theta, alpha and sigma ranges including the dimensions electrode, frequency (4-15 Hz, 1 Hz frequency steps) and time (0-1000ms, five time windows at 200ms each in the WAKE condition, 6 time windows from 0-1200ms during SLEEP). For the “spindle vs. no spindle” and “negative vs. positive SO slope” contrasts we calculated averaged values for FV/UVF for each condition, which we then compared. Please note that for these comparisons we randomly selected a subset of trials so each participant contributed the same number of trials to each of the two conditions to be compared (i.e. for example the same number of “spindle” vs. “no spindle” trials). For all permutation tests the critical *p*-value for the *T*-statistic for dependent samples was set to 0.05 and 1000 randomisations were used. Spatial clusters were formed only if electrodes had a minimum of two neighbouring electrodes that were also significant. We report the Monte Carlo approximation for the estimate of *p*-values. Effects with (one-sided) *Monte Carlo p* < .05 are denoted significant, effects with *Monte Carlo p* < .1 are denoted trends. We report ξ (“explanatory measure of the effect size”) as a robust effect size measure for the comparison of two samples using trimmed means (Wilcox & Tian, 2011), which has been implemented for dependent samples in the ‘yuend’ function in the ‘WSR2’ R package (Mair et al., 2017; Wilcox, 2011). The interpretation of ξ corresponds to Cohen’s d with ξ = .1, .3 and .5 indicating small, medium and large effects, respectively. Critical *p*-values for post-hoc tests were adjusted for multiple comparisons using Bonferroni-Holm-corrected *p*-values. For more details on the statistical analyses please see the supplementary material.

## 3. Results

### 3.1. Wakefulness

Analyses in the delta band (1-3 Hz) yielded a significant effect of *name* (see Fig. 2A and Suppl. Fig. 6). Specifically, analyses revealed that SONs led to stronger ERS at 2 Hz in a frontocentral and a parieto-occipital clusters (*Monte Carlo p* = .01 and *Monte Carlo p* = .02, respectively). This effect was also visible in the ERP with SONs giving rise to a stronger P3 component than UNs (see Fig. 2B). Analyses did not indicate a significant effect of *voice* or a *name* × *voice* interaction (*voice: Monte Carlo ps* > 0.37; *name* × *voice: Monte Carlo ps* > .35). For a summary of all results also see Suppl. Table 1.

**Fig. 2:**
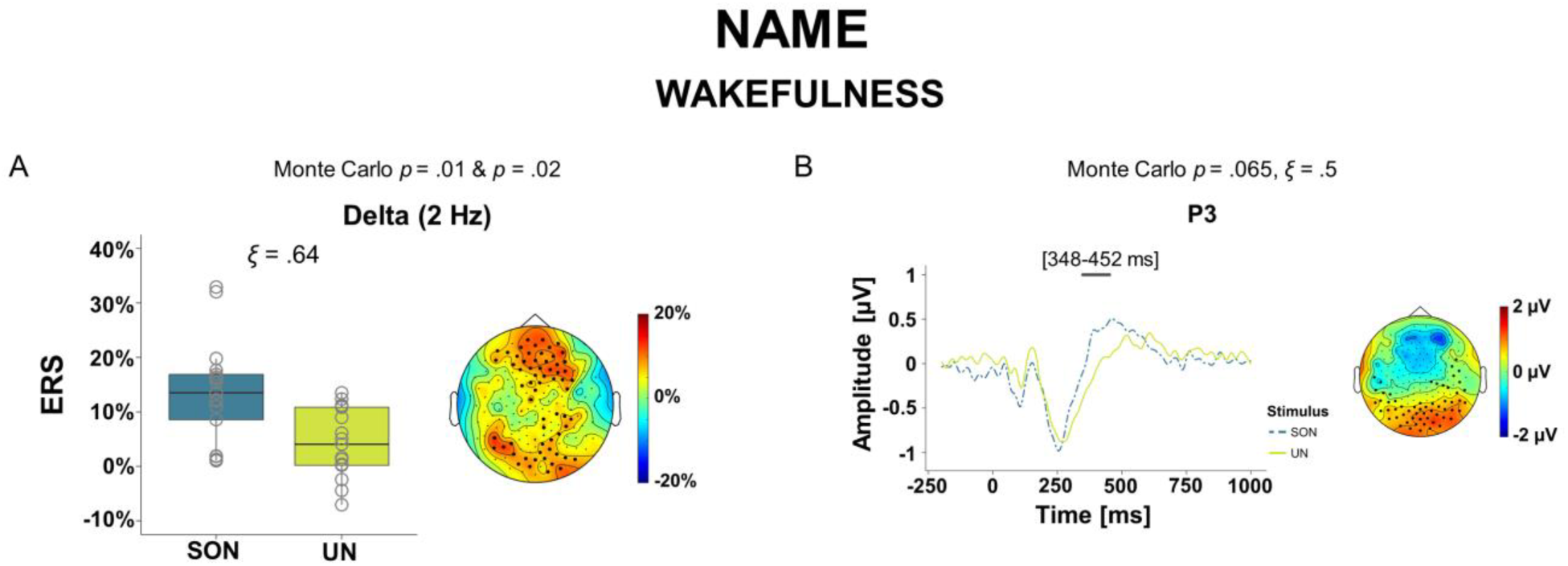
Event-Related Responses during Wakefulness. **(A)** Responses in the delta (1-3 Hz) range. Box plots for the effect of *name* (left) and corresponding scalp plot of differences in ERS between SONs and UNs (right). In box plots, the bold horizontal line corresponds to the median, the lower and upper hinges correspond to the 25^th^ and 75^th^ percentile and the whiskers extend to the lowest/highest values within 1.5 × the interquartile ranges. Open grey circles indicate individual participants’ values. Large black dots indicate the electrodes that are part of the significant clusters at 2 Hz. We report ξ as an estimate of the effect size, with .1, .3 and .5 denoting small, medium and large effects, respectively. Please note that for illustration purposes we show the effects at a representative frequency (i.e. 2 Hz) although significant clusters may have comprised a larger frequency range (see main text and Suppl. Fig. 6). **(B)** Event-related P3 response. Left: Grand average of the ERP elicited by SONs and UNs during wakefulness at all electrodes that were part of the cluster (see scalp plot). The horizontal grey line represents the time window during which the effect was significant (348 to 452ms). Right: Scalp plot of the difference in the ERPs evoked by SONs and UNs. Large black dots indicate electrodes that were part of the cluster with a trend to significance. SON = subject’s own name, UN = unfamiliar name. Analyses and figures are based on data from *n* = 17 participants.

In the theta, alpha and sigma bands (4-15 Hz), the analyses yielded no significant effects (*voice: Monte Carlo ps* > 0.62; *name: Monte Carlo ps* > 0.24; *name* × *voice: Monte Carlo ps* > 0.4).

### 3.2. Sleep

Analysis of the sleep staging results revealed that the median of the total sleep time (TST) during the experimental night was 430.5 minutes (range 300-481.5 min). Wakefulness after sleep onset (WASO) had a median of 20 minutes (range 3.5-110 min). The total number of awakenings varied between 5 and 25 with a median of 15. SOL to N2 was characterised by a median of 20 minutes (range 10-107.5 min), and SOL to REM had a median of 92.5 minutes (range 68.5-228 min). Regarding sleep architecture participants had a median of 7.2% N1 sleep (range 2.7-13.7%), a median of 37% N2 sleep (range 23-54.4%), a median of 34.2% N3 sleep (range 16.5-46.1%) and a median of 18.9% REM sleep (range 12.2-45.1%).

#### 3.2.1. “Enduring Brain State” Analyses

##### 3.2.1.1. N1 sleep

During light N1 sleep, analyses of data from all 17 participants yielded a significant main effect of *voice* in delta (1-3 Hz) ERS (*Monte Carlo p* < .001). Here, UFV stimuli elicited stronger delta ERS than FV stimuli in a cluster that spanned large areas of the scalp with a frontal-central focus (see Fig. 3A and Suppl. Fig. 7A). There were no further significant stimulus-induced differences in the delta range (*name: p* > .21; *name* × *voice:* no significant clusters). Analyses of responses in the theta, alpha and sigma bands (4-15 Hz) also yielded a significant effect of *voice* (*Monte Carlo p* = .003; see Fig. 3C and Suppl. Fig. 7D). Here, UFV stimuli elicited considerable ERS in the theta through sigma frequency range in all time windows analysed (T1-T6: 4-15 Hz). Analyses did not show a significant effect of *name* (*Monte Carlo ps* > .26), but a trend towards a *name* × *voice* interaction (*Monte Carlo p* = .055) with UFV stimuli eliciting stronger ERS irrespective of the name that was presented, i.e. SON or UN, thus confirming the main effect of *voice*. The effects of *voice* were also confirmed by ERP analyses with a stronger positive (92-428ms) and negative component (440-996ms) for UFV as compared to FV (see Suppl. Fig. 2A).

**Fig. 3:**
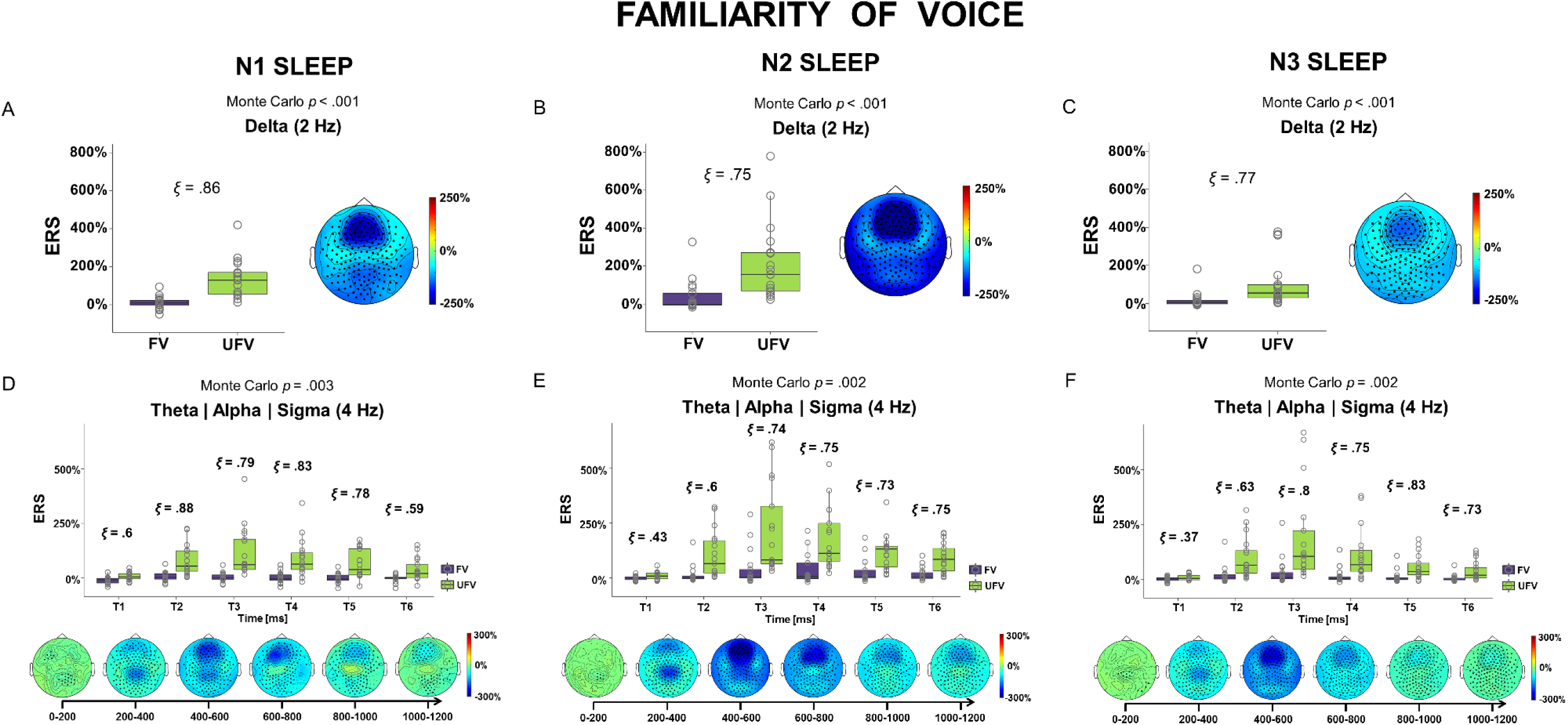
Event-related responses during NREM sleep. **(A, B, C)**: Event-related responses in the delta range (1-3 Hz) during N1, N2 and N3. Box plots for the effect of *voice* (left) and corresponding scalp plots of differences in ERS between FV and UFV (right). **(D, E, F)** Event-related responses in the theta/alpha/sigma range (4-15 Hz) during N1, N2 and N3. Box plots for the effect of *voice* during the six time windows (top) and corresponding scalp plots of differences in ERS/ERD between FV and UFV stimuli (bottom). In box plots, the bold horizontal line corresponds to the median, the lower and upper hinges correspond to the 25^th^ and 75^th^ percentile and the whiskers extend to the lowest/highest values within 1.5 × the interquartile ranges. The open circles are individual participants’ values. We report ξ as an estimate of the effect size, with .1, .3 and .5 denoting small, medium and large effects, respectively. Large black dots indicate the electrodes that are part of the significant clusters. Please note that for illustration purposes we show the effects at representative frequencies (i.e. 2 and 4 Hz) although significant clusters may have comprised a larger frequency range (see main text and Suppl. Fig. 7). FV = familiar voice, UFV = unfamiliar voice. Analyses and figures are based on data from *n* = 17 participants.

##### 3.2.1.2. N2 sleep

Analyses of data from all 17 participants in the delta range yielded a significant effect of *voice* (*p* < .001) with a cluster covering the whole scalp. Specifically, UFV stimuli elicited stronger delta ERS than FV stimuli for all frequencies between 1 and 3 Hz. The (fronto-central) topography was comparable to the N1 effect of *voice* in the delta range (see Fig. 3A, 2B and Suppl. Fig. 7 A and B). Analyses did not yield an effect of *name* or a *name* × *voice* interaction (*Monte Carlo ps* > .18 and *no clusters*, respectively). In the theta to sigma range (4-15 Hz), analyses also revealed a significant effect of *voice* (*Monte Carlo p* = .002). Here, again UFV stimuli elicited strong ERS between 4 and 15 Hz following about 200ms while FV stimuli elicited much less ERS (T1: 4-7 & 15 Hz; T2-T6: 4-15 Hz). The topography and time course was comparable to the N1 effect (see Fig. 3D, Suppl. Fig. 1D and Suppl. Fig. 7E). Besides this, analyses showed no effect of *name* (*Monte Carlo ps* > .16) and no *name* × *voice* interaction (*Monte Carlo ps* >.34). The effects of *voice* in oscillatory analyses were also confirmed by ERP analyses (see Suppl. Fig. 2B).

##### 3.2.1.3. N3 sleep

During N3 sleep, analyses of data from all 17 participants revealed a significant effect of *voice* (*Monte Carlo p* < .001) in the delta range (1-3 Hz). UFV stimuli gave rise to stronger delta ERS than did FV stimuli in a cluster covering large areas of the scalp. Again, the topography was comparable to the results obtained in N1 and N2 (see Fig. 3A-C and Suppl. Fig. 7A-C). Analyses did not reveal any stimulus-induced differences for the *name* effect (no clusters) or the *name* × *voice* interaction (no clusters) in the delta range. Analyses in the theta to sigma range (4-15 Hz) revealed a significant effect of *voice* (*Monte Carlo p* = .002; T1: 4-9 & 15 Hz; T2-6: 14-15 Hz). Here, UFV stimuli elicited stronger ERS than did FV stimuli, an effect that was especially pronounced between about 200 and 1200ms following stimulus onset in a cluster that spanned more or less the whole scalp. Also here, the time course and topography was comparable to the results obtained during N1 and N2 (cf. Fig. 3D-F and Suppl. Fig. 7D-F). Analyses did not yield any other significant effects (*name: Monte Carlo ps* > .23; *name* × *voice: Monte Carlo ps* > .31). Analyses of ERPs confirmed the effects of *voice* (see Suppl. Fig. 2C).

##### 3.2.1.4. REM sleep

Analyses of REM sleep in all 17 participants yielded a significant effect of *voice* in the delta range (1-3 Hz, *Monte Carlo p* = .018, see Fig. 4A and Suppl. Fig. 8A). As during N1-N3, FV stimuli were associated with stronger delta ERS between 1 and 2 Hz than were UFV. There were no further stimulus-induced differences in delta ERS/ERD (*name: Monte Carlo ps* > .18; *name* × *voice* interaction: *Monte Carlo ps* > .17). Analyses in the theta to sigma range (4-15 Hz) yielded a significant *voice* effect (*Monte Carlo p* = .006, see Fig. 4B and Suppl. Fig. 8B). Here, UFV stimuli elicited stronger ERS than FV stimuli following about 200ms. The effect mainly covered the alpha through sigma range (T1: 5 Hz; T2: 4-15 Hz; T3: 5-6 & 12-15 Hz; T4/5: 8-15 Hz; T6: 9-15 Hz) and was most pronounced at the central and centroparietal electrodes. Generally, effects during REM were much less pronounced and delayed compared to the NREM sleep stages. Analyses did not yield any further significant effects (*name: Monte Carlo ps* > .37; *name* × *voice: Monte Carlo ps* > .32).

**Fig. 4:**
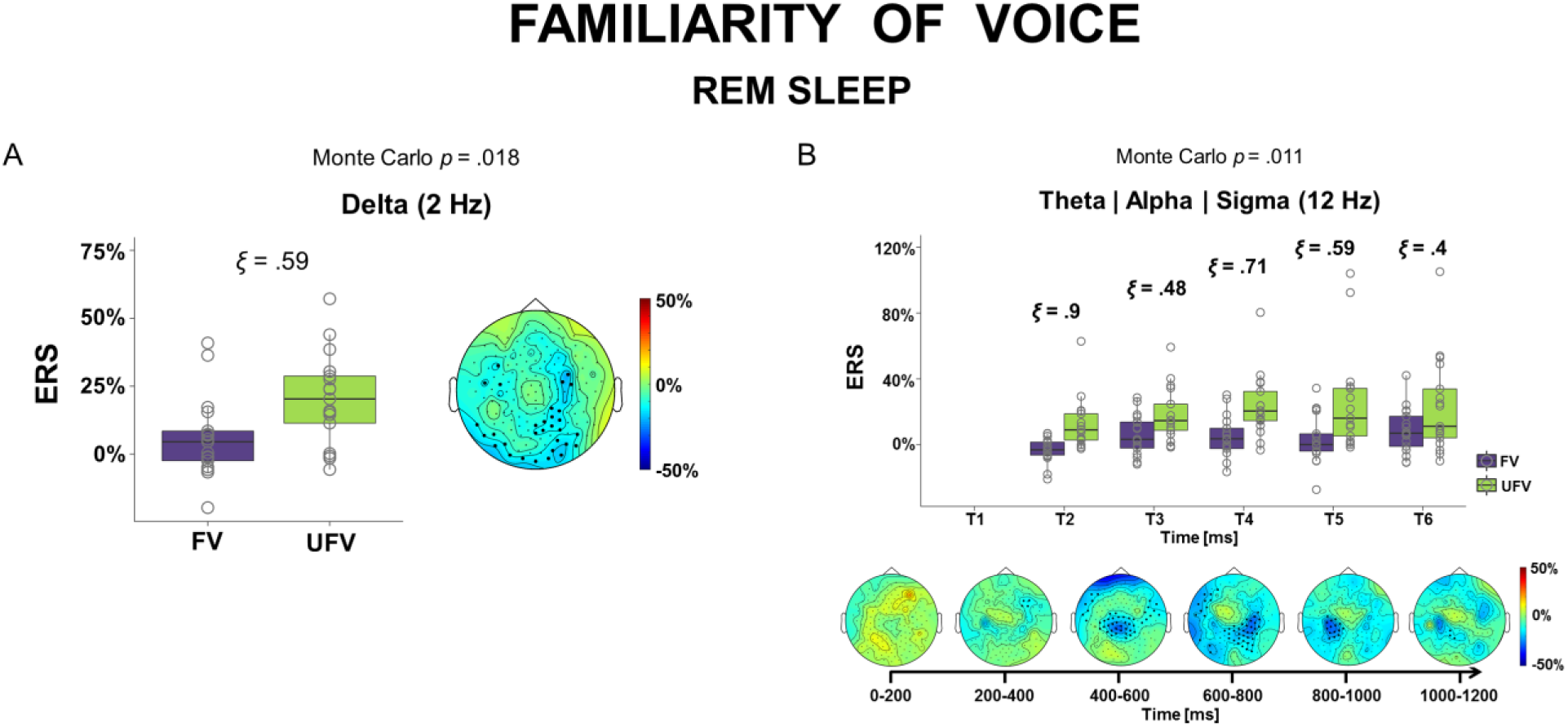
Event-related responses during REM sleep. **(A)** Event-related responses in the delta (1-3 Hz) range. Box plots for the effect of *voice* (left) and corresponding scalp plot of differences in ERS between FV and UFV (right). Large black dots indicate the electrodes that are part of the significant cluster at 2 Hz. **(B)** Event-related responses in the theta/alpha/sigma (4-15 Hz) range. Box plots for the effect of *voice* during the six time windows (top) and corresponding scalp plots of differences in ERS/ERD between FV and UFV stimuli (bottom). In box plots, the bold horizontal line corresponds to the median, the lower and upper hinges correspond to the 25^th^ and 75^th^ percentile and the whiskers extend to the lowest/highest values within 1.5 × the interquartile ranges. Open grey circles indicate individual participants’ values. We report ξ as an estimate of the effect size, with .1, .3 and .5 denoting small, medium and large effects, respectively. Large black dots indicate the electrodes that are part of the cluster at 12 Hz. Please note that for illustration purposes we show the effects at representative frequencies (i.e. 2 and 12 Hz) although significant clusters may have comprised a larger frequency range (see main text and Suppl. Fig. 8). FV = familiar voice, UFV = unfamiliar voice. Analyses and figures are based on data from *n* = 17 participants.

#### 3.2.2. “Transient Brain State”- Analyses

##### 3.2.2.1. Sleep Spindle vs. No Spindle

In both conditions, analyses of ERD/ERS of data from *n* = 14 participants revealed significant effects of *voice* in the delta range (“spindle” condition [S+]: *Monte Carlo p* = .005; 1-3 Hz, see Fig. 4A and Suppl. Fig. 10A and “no spindle” condition [S-]: *Monte Carlo p* = .006; 1-3 Hz, cf. Fig. 4B and Suppl. Fig. 10B) with UFV stimuli eliciting stronger delta ERS than FV stimuli. Post hoc analyses indicated that in the delta range stimulus presentation did not elicit more ERS in the S-compared to the S+ condition (*Monte Carlo ps* > .11). In the S-condition there was also a significant effect of name in the delta range (*Monte Carlo p* = .024, 1-3 Hz) with unfamiliar names (UNs) eliciting stronger ERS than the participant’s own name (SON). There were no further effects in the delta range in either condition (S+: *name: Monte Carlo p* = .14; *name* × *voice: Monte Carlo p* > .22; S-: *name* × *voice: Monte Carlo p* > .49). In the theta to sigma range (4-15 Hz) there were also significant effects of *voice* in both the “spindle” and the “no spindle” conditions (S+: *Monte Carlo p* = .005, T1: 4-7 Hz; T2/3: 4-14 Hz; T4: 4-13 Hz; T5: 4-14 Hz, T6: 4-15 Hz, see Figs. 4C and E and Suppl. Fig. 10C; S-: *Monte Carlo p* = .007, T1: 4 Hz; T2-6: 4-15 Hz, see Figs. 4D and F and Suppl. Fig. 10D). Interestingly, the topography and time course of the effects in the “no spindle” condition were only comparable to the results in the “spindle” condition in the slower frequencies up to about 9 Hz. While in the slower frequencies UFV stimuli elicited stronger ERS than FV stimuli, in the faster frequencies (10-15 Hz), FV stimuli were specifically associated with a marked ERD in the “spindle” condition only (condition differences: *Monte Carlo p* < .001, T1/2: not part of the cluster, T3: 10-15 Hz, T4/5: 9-15 Hz, T6: 8-15 Hz; diamonds in Figs. 4E and F indicate time windows where stimulus-evoked responses were stronger in the S-condition). Analyses did not yield any further significant differences in the theta to sigma range (S+: *name: Monte Carlo ps* > .7 and *name* × *voice: Monte Carlo ps* > .39; S-: *name: Monte Carlo ps* > .32; *name* × *voice: Monte Carlo ps* > .49). ERP analyses showed a significant effect of *voice* that corresponded to the effects in the oscillatory analyses only in the “spindle” condition (see Suppl. Fig. 3).

##### 3.2.2.2. Stimulus Presentation along Slow Oscillation Positive vs. Negative Slope

Irrespective of the slope of a SO during which a stimulus was presented, analyses of data from *n* = 17 participants yielded significant effects of *voice* (pos. slope: *Monte Carlo p* = .001; see Fig. 6A and Suppl. Fig. 11A, neg. slope: *Monte Carlo p* = .002, see Fig. 6B and Suppl. Fig. 11B) in the delta range. Specifically, like in the other conditions UFV stimuli elicited stronger ERS than did FV stimuli between 1 and 3 Hz in clusters spanning large parts of the scalp. However, in the positive SO slope condition stimulus presentation and in particular UFV stimuli elicited significantly larger responses than in the negative SO slope condition at 1 Hz (*Monte Carlo p* < .02). No further effects were evident in the delta range (pos. slope: *name: Monte Carlo p* = .11; *voice × name:* no clusters; neg. slope: *name: Monte Carlo p* = .15; *voice × name: Monte Carlo p* > .17). In the theta to sigma range (4-15 Hz) analyses also revealed significant effects of *voice* in both conditions (pos. slope: *Monte Carlo p* < .001, see Fig. 6C and Suppl. Fig. 11C; neg. slope: *Monte Carlo p* = .004, see Fig. 6D and Suppl. Fig. 11D) with UFV stimuli eliciting stronger ERS than FV stimuli following about 200ms in a broad frequency range comparable to the effects in the other conditions regarding topography and time course (pos. slope: T1: 13-15 Hz; T2-6: 4-15 Hz; neg. slope: T1: 4-11 Hz; T2-6: 4-15 Hz). There were significant differences between the positive and negative SO slope conditions (*Monte Carlo ps* < .009) with stimulus presentation eliciting stronger responses in the positive SO slope condition beyond about 200ms. The effects of *voice* were also confirmed by ERP analyses (pos. slope: see Suppl. Fig. 4B; neg. slope: see Suppl. Fig. 4C). There were no further effects in the theta through sigma range (pos. slope: *name: Monte Carlo ps* > .30; *name* × *voice: Monte Carlo ps* > .28; neg. slope: *name: Monte Carlo ps* > .20; *name × voice: Monte Carlo ps* >.12).

**Fig. 6:**
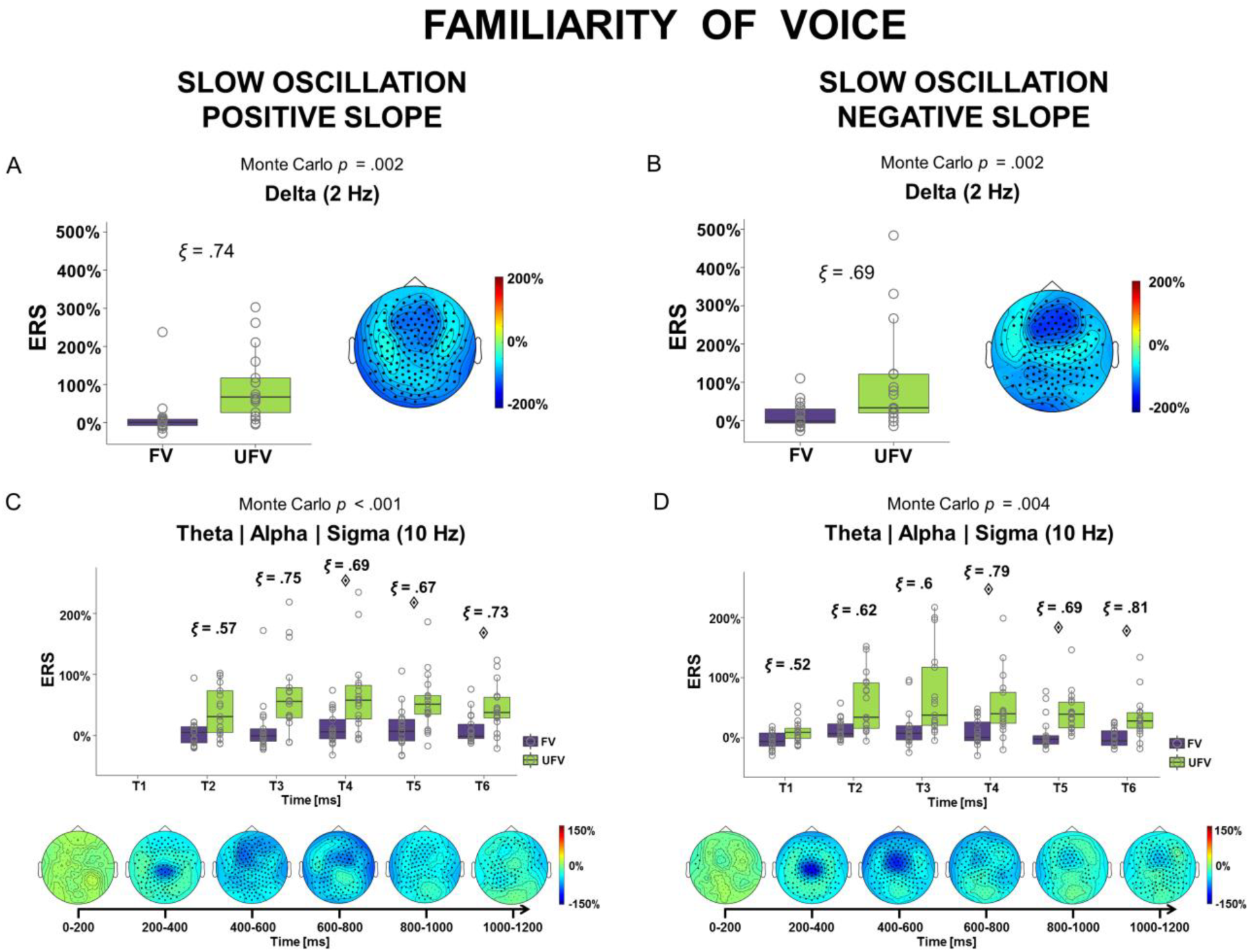
Event-related responses during N2/N3 sleep when a stimulus was presented along the positive vs. negative slope of the SO. **(A, B)** Event-related responses in the delta (1-3 Hz) range. Box plots for the effect of *voice* (left) and corresponding scalp plots of differences in ERS between FV and UFV (right). **(C, D)** Event-related responses in the theta/alpha/sigma (4-15 Hz) range. Box plots for the effect of *voice* during the six time windows (top) and corresponding scalp plots of differences in ERS/ERD between FV and UFV stimuli (bottom). In box plots, the bold horizontal line corresponds to the median, the lower and upper hinges correspond to the 25^th^ and 75^th^ percentile and the whiskers extend to the lowest/highest values within 1.5 × the interquartile ranges. Open grey circles indicate individual participants’ values. We report ξ as an estimate of the effect size, with .1, .3 and .5 denoting small, medium and large effects, respectively. Diamonds indicate time windows during which stimulus-induced differences were more pronounced in the positive SO slope condition. Large black dots indicate the electrodes that are part of the significant clusters. Please note that for illustration purposes we show the effects at representative frequencies (i.e. 2 and 10 Hz) although significant clusters may have comprised a larger frequency range (see main text and Suppl. Fig. 11). FV = familiar voice, UFV = unfamiliar voice. Analyses and figures are based on data from *n* = 17 participants.

## 4. Discussion

In this study we show that especially processing of paralinguistic emotional aspects of verbal stimuli such as the familiarity of a voice is even possible during fading and in the absence of consciousness defined as behavioural responsiveness during sleep. The findings add to existing evidence that the detection and evaluation of meaningful stimuli is still possible in these states (e.g. Perrin et al., 1999; Portas et al., 2000). Intriguingly, we do not only find that a differential response to familiar vs. unfamiliar voice (FV vs. UFV) stimuli persists during light NREM sleep stages N1 and N2 thus replicating previous results (cf. Blume et al., 2016; Perrin et al., 1999), but we extend this finding to deep N3 and also REM sleep. Beyond this, we show that transient brain states, which have been suggested to alter sensory information processing during sleep, i.e. sleep spindles (Cote et al., 2000; Elton et al., 1997; Schabus et al., 2012) and slow oscillation down-states (Massimini et al., 2003; Schabus et al., 2012), do, at least not uniformly or irrespective of stimulus characteristics, inhibit stimulus processing. Rather, their inhibitory function seems to be tuned to stimulus salience.

During wakefulness, SONs seemed to be salient when compared to UNs thus drawing more attentional resources. This was indicated by SONs eliciting stronger delta ERS than UNs across large areas of the scalp. Functionally, delta ERS has repeatedly been linked to attentional processes and the detection of salient or motivationally relevant stimuli (for reviews see Knyazev, 2007; Knyazev, 2012). Additionally, this is well in line with the relatively larger P3 component evident in ERP analyses, which has likewise been associated with attention and stronger processing, as well as results from earlier studies (e.g. Berlad & Pratt, 1995; Blume et al., 2016; Perrin et al., 1999). In an earlier study from our group, del Giudice et al. (2014) had also found stronger alpha ERD for SONs than for UNs, which we could not replicate here. This may ultimately be due to methodological differences and the more conservative statistical analysis methods employed here. Somewhat surprisingly, no differences were evident between FV and UFV stimuli when participants were awake, although earlier studies had reported differential effects on ERPs (Beauchemin et al., 2006; Holeckova et al., 2006). Also here, methodological differences may account for the deviating findings. Nevertheless it should be noted that we experience situations in which voices of varying degrees of familiarity are present along with unfamiliar voices every day. From this perspective, one may speculate whether the lack of a differential response evoked by voice familiarity may even indicate adaptive processing mechanisms precluding the mere presence of familiar voices from interfering with targeted attentional processes.

During NREM sleep, that is from light N1 to deep N3 sleep, we consistently find that processing of FV vs. UFV stimuli gives rise to a differential response in the delta to sigma frequency range, an effect that is present in oscillatory analyses as well as ERPs. Most importantly, this provides support for the notion that processing of auditory stimuli and especially of paralinguistic stimulus aspects such as the familiarity of a voice is incessantly processed even in states where consciousness is absent. While this is well in line with earlier findings during light sleep stages N1 and N2 (e.g. Blume et al., 2016; Oswald et al., 1960; Portas et al., 2000), our results suggest that the same holds true even for deep N3 sleep. Thus, the findings also support the notion of a ‘sentinel processing mode’ of the brain during sleep, which we suggested in a previous publication (cf. Blume et al., 2016). Specifically, this mode describes the idea that (low-level) stimulus evaluation continues even when consciousness fades during sleep and the result of this evaluation may subsequently either trigger an inhibitory sleep-protecting response or awakening. In detail, we here find UFV stimuli to be associated with stronger ERS in the delta range than FV stimuli during all NREM sleep stages, an effect which was widespread across the scalp with the response being most pronounced above frontocentral areas. Adopting the interpretation of delta oscillations during wakefulness, the results suggest that UFV stimuli may become salient when consciousness fades (Knyazev, 2007, 2012). In particular, the presence of unfamiliar voices could challenge the impression of a safe environment that is necessary to ‘let go of consciousness’ and eventually fall and stay asleep, rendering them salient. However, the increase in delta ERS visible in oscillatory analyses could also reflect a sleep-specific event-related pattern, namely an evoked slow oscillatory or K-complex-like response. Like slow oscillations (SOs), K-complexes (KCs) have their peak frequency is in the delta range and they are considered ‘forerunners’ of SOs or ‘sub-threshold SOs’ (e.g. Amzica & Steriade, 1997; De Gennaro et al., 2000), which are often evoked by acoustic stimulation (Bellesi et al., 2014). While KCs are strongly associated with N2 sleep, SOs are considered the hallmark of N3 sleep. Functionally, evoked slow waves (i.e. KCs and SOs), have been suggested to serve cortical excitation and low-level information processing as well as the subsequent protection of sleep by neuronal silencing (Cash et al., 2009; Dang-Vu et al., 2011; Laurino et al., 2014). Although they also occur spontaneously, especially KCs have been found to be elicited particularly by salient or high-intensity stimuli (e.g. Bastien & Campbell, 1992). In line with the notion that evoked slow waves indicate ongoing cognitive processing, Vallat et al. (2017) have recently reported a KC/SO-like response during N2 sleep that was stronger for auditory stimuli that were followed by an arousal or awakening. The authors concluded that this reflects stronger reactivity of the brain to external stimuli, which in turn leads to stronger arousal. In accordance with this, ERP analyses of our data indicated that stimulus-induced differences in the delta range indeed reflected KC/SO-like responses evoked by stimulus presentation with considerably larger amplitudes for UFV stimuli. In line with earlier ideas, we suggest that this ERP reflects increased (low-level) information processing of especially salient UFV stimuli (indexed by a larger positive wave), which is then followed by an inhibitory or sleep-protecting ‘down-state’ (indexed by a larger negative wave) that is likewise scaled to stimulus salience. Further support for this interpretation comes from analyses when we explicitly looked at stimulus presentations that evoked an SO, with evoked SOs also seeming to be sensitive to stimulus salience. Also here, UFV stimuli were associated with stronger delta through sigma activity than FV stimuli and ERP analyses revealed that UFV stimuli were associated with a very slight positive-going wave, which was followed by a SO down-state that appeared much more pronounced for UFV stimuli (cf. Suppl. Fig. 4A). Besides the results obtained in the delta range, we also find that during all NREM sleep stages UFV stimuli are associated with stronger ERS in the theta through sigma range than FV stimuli, an effect which is most pronounced following about 200ms after stimulus onset. Most importantly, these findings are well in line with the delta results and they provide further convincing support for the notion that the brain is still able to process paralinguistic stimulus aspects even when consciousness fades and is absent. On a functional level, especially frequencies in the alpha and sigma range are thought to mirror an increase in arousal during sleep (cf. American Academy of Sleep Medicine & Iber, 2007). This suggests that UFV stimuli may be more arousing than FV stimuli during NREM sleep, an interpretation that, also given the observed KC/SO-like response, is well in line with Vallat et al.’s results. As suggested above, the presence of unfamiliar voices may challenge the impression of an environment ‘safe to sleep’ and thus be arousing. Admittedly, our findings during N2 sleep partly contrast results of earlier studies, where the brain also seemed to continue differentiating between UNs and SONs (e.g. Blume et al., 2016; Perrin et al., 1999). The deviating findings are likely to be due to methodological differences. Additionally, it should be noted that in the present study participants slept during a whole night and not just an afternoon nap (cf. Blume et al., 2016) with differences in the homeostatic and circadian factors rendering it questionable whether a daytime nap can be considered a short night sleep equivalent (Dijk & Czeisler, 1995; van Schalkwijk et al., 2017).

In summary, results obtained during wakefulness and NREM sleep suggest that familiarity of a voice can be processed even during the fading of consciousness (N1) and in the full absence of (behavioural) consciousness (N2 and N3). For REM sleep, a paradoxical state characterised by (i) the return of ‘altered consciousness’, namely ‘dreaming’ (although note that dreams are not limited to REM sleep, cf. e.g. Siclari et al., 2017), (ii) enhanced brain metabolism compared to wakefulness (Nofzinger et al., 1997) and (iii) an increase in higher frequency EEG power (Uchida et al., 1992), we also observed a relatively stronger increase in delta as well as alpha/sigma ERS elicited by UFV compared to FV stimuli, which may indicate continued processing and/or arousal of salient or potentially ‘dangerous’ UFV stimuli. This is especially interesting because REM sleep has been suggested to reflect a ‘closed loop’, that is a state in which the brain is rather occupied with intrinsic activity than processing of external stimuli (Andrillon et al., 2016; Llinás & Paré, 1991; Wehrle et al., 2007) with our results challenging this notion. At the same time, while the oscillatory response pattern was generally similar to NREM sleep findings, REM responses were considerably weaker (see also Suppl. Fig. 9) and markedly delayed by approx. 400ms. This underlines the idea that brain activity and processing of environmental stimuli during REM is qualitatively different although not generally precluded. In conclusion, we consistently find that during all NREM sleep stages as well as REM sleep, the brain seems to continue differentiating between paralinguistic (emotional) aspects (i.e. familiar vs. unfamiliar voice) but not among the linguistic content of stimuli (i.e. own vs. other name; cf. Suppl. Table 1 for an overview of the results). In contrast to processing of the content, which involves higher level cognitive processes including for example memory access, processing of emotional content and the identity of a voice has been suggested to be possible also at lower levels. It has for example been reported that the identification of emotions or identity in voice occurs at very early stages of processing (emotions at about 200ms, identity at about 300ms already; cf. Spreckelmeyer et al., 2009) and emotional prosody processing occurs in regions close to primary auditory regions and irrespective of the listeners’ focus of attention (Grandjean et al., 2005). From this perspective, it seems that during sleep, which is characterised by the reduced availability of cognitive resources, the brain may be apt to processing of paralinguistic (emotional) stimulus characteristics.

Beyond investigating stimulus processing across enduring brain states, i.e. wakefulness and different sleep stages, we were also interested in how stimulus presentation relates to ‘transient oscillatory activity’, that is sleep spindles and slow oscillations (SOs), during N2 and N3 sleep. Generally, sleep spindles (Elton et al., 1997; Schabus et al., 2012) and the negative slope of slow oscillations (Schabus et al., 2012) have been suggested to inhibit processing of external stimuli. In line with this we find that sleep spindles as well as a negative slow oscillation slope attenuate stimulus processing (cf. Fig. 5E/F and Suppl. Fig. 4 B/C). However, this does not seem to be an all-or-none phenomenon, but rather brain responses are still tuned to stimulus salience suggesting that at least ‘low-level’ processing is not precluded. More specifically, we find that when a sleep spindle overlapped with stimulus presentation UFV stimuli still elicited responses in the delta through lower alpha (i.e. up to about 9 Hz) range that were similar to those obtained when not taking ‘transient oscillatory activity’ into account. Intriguingly and unlike proposed earlier (Schabus et al., 2012; Steriade, 1991), this suggests that processing of external stimuli is not or at least not uniformly inhibited by the presence of a sleep spindle, i.e. spindles do not generally seem to act as a sensory filter at the thalamic level. Interestingly, this is well in line with recent findings in rodents where thalamocortical sensory relay was shown to persist even during sleep spindles (Sela et al., 2016). Beyond this, above ≈9 Hz the response pattern when a spindle was present was markedly different from the general NREM (see Fig. 3) and, most importantly, the ‘no spindle’ (see Fig. 5F) patterns with FV stimuli eliciting stronger ERD than UFV stimuli in the ≈11-15 Hz spindle range (see Fig. 5C). We speculate that this could reflect a relatively stronger release of inhibition (reflected by 10-15 Hz ERD) for seemingly less relevant FV stimuli by sleep spindles. Arguably, a selective mechanism that specifically filters information that is considered irrelevant, i.e. here FV stimuli, seems more adaptive than the uniform inhibition of all environmental stimuli. Following the idea of a ‘sentinel processing mode’ of the brain during sleep, spindles just as slow oscillations could thus reflect a sleep-protecting response that follows initial stimulus evaluation during N2 and N3. Besides sleep spindles, previous studies suggested that also the slope of a SO during stimulus presentation affects stimulus processing. In particular the negative slope has been found to be associated with decreased responses in studies using somatosensory stimuli and simple tones as compared to the positive SO slope (Dang-Vu et al., 2011; Massimini et al., 2003; Schabus et al., 2012). Surprisingly, in our study stimulus delivery during negative and positive slopes revealed similar responses with responses in both conditions being tuned to stimulus salience. Specifically, as during all other sleep stages UFV stimuli elicited stronger (delta to sigma) ERS than FV stimuli. These results were supported by ERP analyses indicating that UFV stimuli induced a more pronounced down-state that was preceded by an up-state. The findings thereby contrast earlier findings and suggest that also the negative slope of a SO does at least not uniformly inhibit information processing and allows continued evaluation of stimulus characteristics. Likewise, the findings also suggest that during a positive SO slope the brain is not uniformly open to external stimulation.

**Fig. 5:**
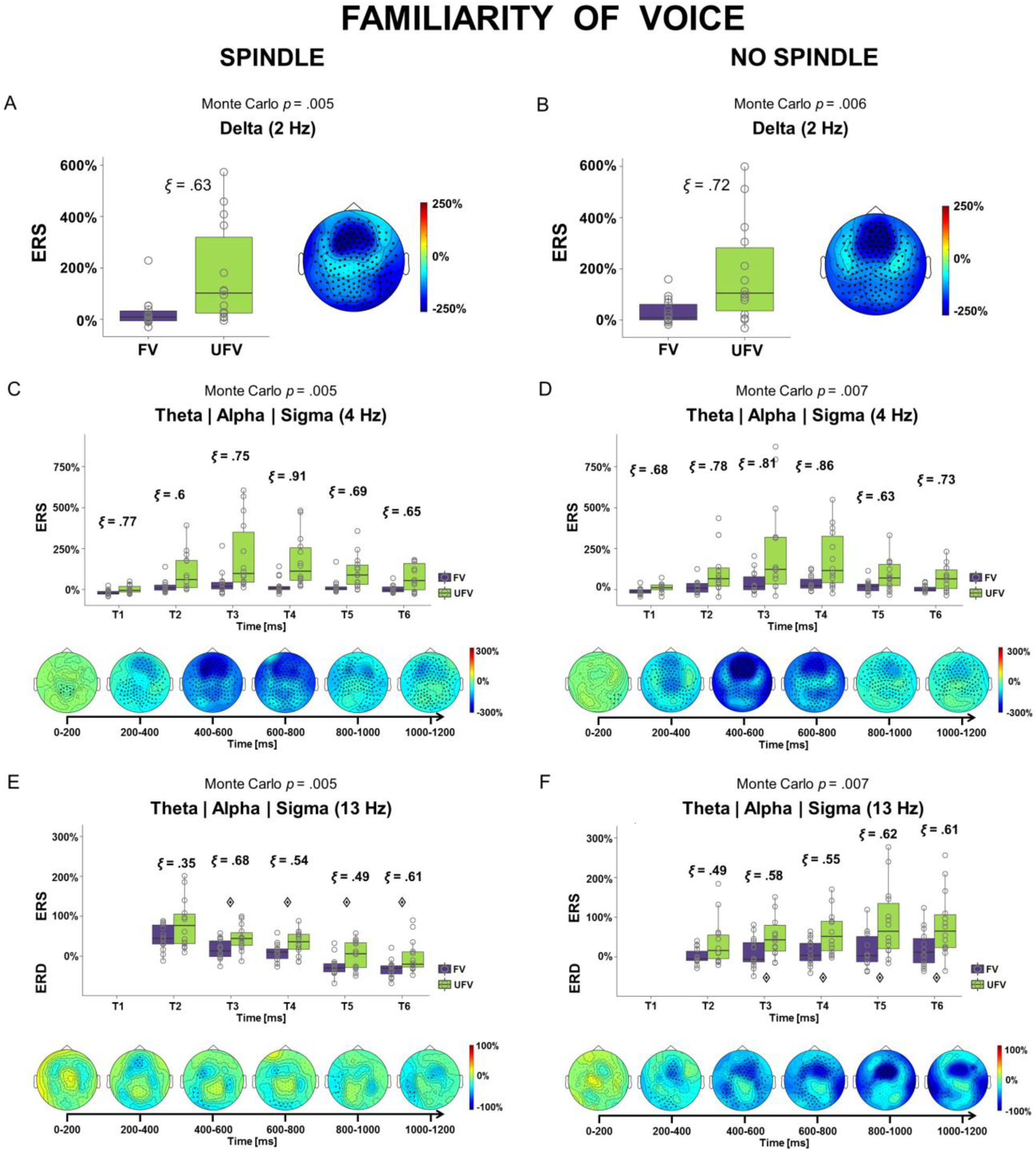
Event-related responses during N2/N3 sleep depending on the presence/absence of sleep spindles. **(A, B)** Event-related responses in the delta (1-3 Hz) range. Box plots for the effect of *voice* (left) and corresponding scalp plot of differences in ERS between FV and UFV (right). **(C, D)** Event-related responses in the theta/alpha/sigma (4-15 Hz) range at 4 Hz and **(E, F)** responses at 13 Hz. Box plots for the effect of *voice* during the six time windows (top) and corresponding scalp plots of differences in ERS/ERD between FV and UFV stimuli (bottom). In box plots, the bold horizontal line corresponds to the median, the lower and upper hinges correspond to the 25^th^ and 75^th^ percentile and the whiskers extend to the lowest/highest values within 1.5 × the interquartile ranges. Open grey circles indicate individual participants’ values. Diamonds in figures E and F indicate the time windows with significant differences between S+ and S-conditions at 13 Hz. We report ξ as an estimate of the effect size, with .1, .3 and .5 denoting small, medium and large effects, respectively. Large black dots indicate the electrodes that are part of the clusters at 2 Hz, 4 Hz or 13 Hz, respectively. Please note that for illustration purposes we show the effects at representative frequencies (i.e. 2, 4 and 13 Hz) although significant clusters may have comprised a larger frequency range (see main text and Suppl. Fig. 10). A spindle could either be present during stimulus onset (i.e. spindle offset min. 200ms after stimulus onset) or it could have a substantial overlap with stimulus presentation (spindle onset 0-400ms after stimulus onset). For more details please see supplementary material. FV = familiar voice, UFV = unfamiliar voice. Analyses and figures are based on data from *n* = 14 participants.

In conclusion, this study shows that stimulus characteristics and especially the familiarity of a voice continue to be evaluated during all stages of NREM sleep and thus even in the complete absence of behavioural consciousness. Surprisingly, this is the case even during REM sleep with processing of external seeming to be slowed and decreased though. Our findings thereby provide support for the idea of a ‘sentinel processing mode’ of the brain during sleep, i.e. the continued processing of environmental stimuli even in the absence of consciousness that may then be followed by either an inhibitory sleep-protective response or awakening depending on the result of stimulus evaluation. Beyond this, it appears that even ‘transient oscillatory activity’, i.e. sleep spindles and slow oscillations are sensitive to paralinguistic emotional stimulus characteristics. Furthermore, we provide novel evidence that, although stimulus processing is generally attenuated, even during spindles and the negative slope of a SO the brain reacts differentially to incoming information. More generally, our findings also suggest that in different vigilance states processing of emotional stimuli may vary. Besides, the results may open up new perspectives for insomnia research, where a relative deficit in processing of environmental stimuli during sleep may be related to problems of ‘letting go of consciousness’ and thus falling asleep.

## Acknowledgements

We thank Daniel Koerner and Adriana Michalak for their help with the data collection and pre-processing and Vincenzo Muto for his advice on the sleep staging. We also thank Wolfgang Klimesch for the valuable discussion of the results.

## Supporting Information

A: Supplementary Material (Methods, Results)

## References

American Academy of Sleep Medicine, & Iber, C. (2007). The AASM manual for the scoring of sleep and associated events: rules, terminology and technical specifications: American Academy of Sleep Medicine.

Amzica, F., & Steriade, M. (1997). The K-complex: Its slow (<1-Hz) rhythmicity and relation to delta waves. Neurology, 49(4), 952–959. doi:10.1212/wnl.49.4.952

Anderer, P., Gruber, G., Parapatics, S., Woertz, M., Miazhynskaia, T., Klösch, G., … Danker-Hopfe, H. (2005). An E-health solution for automatic sleep classification according to Rechtschaffen and Kales: validation study of the Somnolyzer 24× 7 utilizing the Siesta database. Neuropsychobiology, 51(3), 115–133.

Anderer, P., Moreau, A., Woertz, M., Ross, M., Gruber, G., Parapatics, S., … Dorffner, G. (2010). Computer-Assisted Sleep Classification according to the Standard of the American Academy of Sleep Medicine: Validation Study of the AASM Version of the Somnolyzer 24 × 7. Neuropsychobiology, 62(4), 250–264.

Anderer, P., Saletu, B., Saletu-Zyhlarz, G. M., Gruber, G., Parapatics, S., Miazhynskaia, T., … Dorffner, G. (2004). Recent advances in the electrophysiological evaluation of sleep. Essentials and applications of EEG research in preclinical and clinical pharmacology. Berlin: Unipublish Verlag für Studium & Praxis OHG, 307–339.

Andrillon, T., Poulsen, A. T., Hansen, L. K., Léger, D., & Kouider, S. (2016). Neural Markers of Responsiveness to the Environment in Human Sleep. The Journal of Neuroscience, 36(24), 6583–6596. doi:10.1523/jneurosci.0902-16.2016

Bastien, C., & Campbell, K. (1992). The evoked K-complex: all-or-none phenomenon? Sleep, 15(3), 236–245.

Bastuji, H., & García-Larrea, L. (1999). Evoked potentials as a tool for the investigation of human sleep. Sleep medicine reviews, 3(1), 23–45. doi:http://dx.doi.org/10.1016/S1087-0792(99)90012-6

Beauchemin, M., Beaumont, L. D., Vannasing, P., Turcotte, A., Arcand, C., Belin, P., & Lassonde, M. (2006). Electrophysiological markers of voice familiarity. European Journal of Neuroscience, 23(11), 3081–3086. doi:doi:10.1111/j.1460-9568.2006.04856.x

Bellesi, M., Riedner, B. A., Garcia-Molina, G. N., Cirelli, C., & Tononi, G. (2014). Enhancement of sleep slow waves: underlying mechanisms and practical consequences. Frontiers in Systems Neuroscience, 8, 208. doi:10.3389/fnsys.2014.00208

Berlad, I., & Pratt, H. (1995). P300 in response to the subject’s own name. Electroencephalogr Clin Neurophysiol, 96(5), 472–474.

Blume, C., del Giudice, R., Lechinger, J., Wislowska, M., Heib, D. P. J., Hoedlmoser, K., & Schabus, M. (2016). Preferential processing of emotionally and self-relevant stimuli persists in unconscious N2 sleep. Brain Lang. doi:http://dx.doi.org/10.1016/j.bandl.2016.02.004

Bullmore, E. T., Suckling, J., Overmeyer, S., Rabe-Hesketh, S., Taylor, E., & Brammer, M. J. (1999). Global, voxel, and cluster tests, by theory and permutation, for a difference between two groups of structural MR images of the brain. IEEE Transactions on Medical Imaging, 18(1), 32–42. doi:10.1109/42.750253

Cash, S. S., Halgren, E., Dehghani, N., Rossetti, A. O., Thesen, T., Wang, C., … Madsen, J. R. (2009). The human K-complex represents an isolated cortical down-state. Science, 324(5930), 1084–1087.

Cote, K. A., De Lugt, D. R., Langley, S. D., & Campbell, K. B. (1999). Scalp topography of the auditory evoked K-complex in stage 2 and slow wave sleep. Journal of Sleep Research, 8(4), 263–272.

Cote, K. A., Epps, T. M., & Campbell, K. B. (2000). The role of the spindle in human information processing of high-intensity stimuli during sleep. Journal of Sleep Research, 9(1), 19–26. doi:10.1046/j.1365-2869.2000.00188.x

Dang-Vu, T. T., Bonjean, M., Schabus, M., Boly, M., Darsaud, A., Desseilles, M., … Luxen, A. (2011). Interplay between spontaneous and induced brain activity during human non-rapid eye movement sleep. Proceedings of the National Academy of Sciences, 108(37), 15438–15443.

De Gennaro, L., & Ferrara, M. (2003). Sleep spindles: an overview. Sleep medicine reviews, 7(5), 423–440.

De Gennaro, L., Ferrara, M., & Bertini, M. (2000). The spontaneous K-complex during stage 2 sleep: is it the ‘forerunner’ of delta waves? Neuroscience Letters, 291(1), 41–43. doi:https://doi.org/10.1016/S0304-3940(00)01366-5

del Giudice, R., Lechinger, J., Wislowska, M., Heib, D. P., Hoedlmoser, K., & Schabus, M. (2014). Oscillatory brain responses to own names uttered by unfamiliar and familiar voices. Brain Res, 1591, 63–73.

Dijk, D.-J., & Czeisler, C. A. (1995). Contribution of the circadian pacemaker and the sleep homeostat to sleep propensity, sleep structure, electroencephalographic slow waves, and sleep spindle activity in humans. The Journal of Neuroscience, 15(5), 3526–3538.

Dijk, D.-J., Duffy, J. F., & Czeisler, C. A. (1992). Circadian and sleep/wake dependent aspects of subjective alertness and cognitive performance. Journal of Sleep Research, 1(2), 112–117. doi:10.1111/j.1365-2869.1992.tb00021.x

Elton, M., Winter, O., Heslenfeld, D., Loewy, D., Campbell, K., & Kok, A. (1997). Event-related potentials to tones in the absence and presence of sleep spindles. Journal of Sleep Research, 6(2), 78–83.

Grandjean, D., Sander, D., Pourtois, G., Schwartz, S., Seghier, M. L., Scherer, K. R., & Vuilleumier, P. (2005). The voices of wrath: brain responses to angry prosody in meaningless speech. Nature neuroscience, 8, 145. doi:10.1038/nn1392 https://www.nature.com/articles/nn1392#supplementary-information

Heib, D. P., Hoedlmoser, K., Anderer, P., Zeitlhofer, J., Gruber, G., Klimesch, W., & Schabus, M. (2013). Slow Oscillation Amplitudes and Up-State Lengths Relate to Memory Improvement. PLoS One, 8(12), e82049. doi:10.1371/journal.pone.0082049

Holeckova, I., Fischer, C., Giard, M. H., Delpuech, C., & Morlet, D. (2006). Brain responses to a subject’s own name uttered by a familiar voice. Brain Res, 1082(1), 142–152.

Klimesch, W. (1999). EEG alpha and theta oscillations reflect cognitive and memory performance: a review and analysis. Brain Research Reviews, 29(2-3), 169–195.

Klimesch, W., Doppelmayr, M., Russegger, H., Pachinger, T., & Schwaiger, J. (1998). Induced alpha band power changes in the human EEG and attention. Neuroscience Letters, 244(2), 73–76. doi:http://dx.doi.org/10.1016/S0304-3940(98)00122-0

Klimesch, W., Schack, B., & Sauseng, P. (2005). The functional significance of theta and upper alpha oscillations. Experimental Psychology, 52(2), 99–108.

Knyazev, G. G. (2007). Motivation, emotion, and their inhibitory control mirrored in brain oscillations. Neuroscience & Biobehavioral Reviews, 31(3), 377–395. doi:http://dx.doi.org/10.1016/j.neubiorev.2006.10.004

Knyazev, G. G. (2012). EEG delta oscillations as a correlate of basic homeostatic and motivational processes. Neuroscience & Biobehavioral Reviews, 36(1), 677–695. doi:http://dx.doi.org/10.1016/j.neubiorev.2011.10.002

Laurino, M., Menicucci, D., Piarulli, A., Mastorci, F., Bedini, R., Allegrini, P., & Gemignani, A. (2014). Disentangling different functional roles of evoked K-complex components: Mapping the sleeping brain while quenching sensory processing. Neuroimage, 86, 433–445. doi:http://dx.doi.org/10.1016/j.neuroimage.2013.10.030

Llinás, R. R., & Paré, D. (1991). Of dreaming and wakefulness. Neuroscience, 44(3), 521–535. doi:http://dx.doi.org/10.1016/0306-4522(91)90075-Y

Mair, P., Schoenbrodt, F., & Wilcox, R. R. (2017). WRS2: Wilcox robust estimation and testing (Version 0.9-2).

Maris, E., & Oostenveld, R. (2007). Nonparametric statistical testing of EEG-and MEG-data. Journal of Neuroscience Methods, 164(1), 177–190.

Massimini, M., & Amzica, F. (2001). Extracellular calcium fluctuations and intracellular potentials in the cortex during the slow sleep oscillation. Journal of neurophysiology, 85(3), 1346–1350.

Massimini, M., Rosanova, M., & Mariotti, M. (2003). EEG slow (∼ 1 Hz) waves are associated with nonstationarity of thalamo-cortical sensory processing in the sleeping human. Journal of neurophysiology, 89(3), 1205–1213.

Nofzinger, E. A., Mintun, M. A., Wiseman, M., Kupfer, D. J., & Moore, R. Y. (1997). Forebrain activation in REM sleep: an FDG PET study. Brain Res, 770(1), 192–201.

Oostenveld, R., Fries, P., Maris, E., & Schoffelen, J.-M. (2010). FieldTrip: open source software for advanced analysis of MEG, EEG, and invasive electrophysiological data. Computational intelligence and neuroscience, 2011.

Oswald, I., Taylor, A. M., & Treisman, M. (1960). Discriminative responses to stimulation during human sleep. Brain.

Perrin, F., Garcia-Larrea, L., Mauguiere, F., & Bastuji, H. (1999). A differential brain response to the subject’s own name persists during sleep. Clinical Neurophysiology, 110(12), 2153–2164.

Portas, C. M., Krakow, K., Allen, P., Josephs, O., Armony, J. L., & Frith, C. D. (2000). Auditory processing across the sleep-wake cycle: simultaneous EEG and fMRI monitoring in humans. Neuron, 28(3), 991–999.

Riedner, B. A., Vyazovskiy, V. V., Huber, R., Massimini, M., Esser, S., Murphy, M., & Tononi, G. (2007). Sleep homeostasis and cortical synchronization: III. A high-density EEG study of sleep slow waves in humans. Sleep, 30(12), 1643–1657.

Santhi, N., Lazar, A. S., McCabe, P. J., Lo, J. C., Groeger, J. A., & Dijk, D.-J. (2016). Sex differences in the circadian regulation of sleep and waking cognition in humans. Proc Natl Acad Sci U S A, 113(19), E2730–E2739. doi:10.1073/pnas.1521637113

Schabus, M., Dang-Vu, T. T., Heib, D. P. J., Boly, M., Desseilles, M., Vandewalle, G., … Gais, S. (2012). The fate of incoming stimuli during NREM sleep is determined by spindles and the phase of the slow oscillation. Frontiers in neurology, 3.

Sela, Y., Vyazovskiy, V. V., Cirelli, C., Tononi, G., & Nir, Y. (2016). Responses in rat core auditory cortex are preserved during sleep spindle oscillations. Sleep, 39(5), 1069–1082.

Siclari, F., Baird, B., Perogamvros, L., Bernardi, G., LaRocque, J. J., Riedner, B., … Tononi, G. (2017). The neural correlates of dreaming. Nature neuroscience.

Spreckelmeyer, K. N., Kutas, M., Urbach, T., Altenmüller, E., & Münte, T. F. (2009). Neural processing of vocal emotion and identity. Brain Cogn, 69(1), 121–126. doi:https://doi.org/10.1016/j.bandc.2008.06.003

Steriade, M. (1991). Normal and Altered States of Function. Cerebral Cortex, 9, 279e357.

Strauss, M., Sitt, J. D., King, J.-R., Elbaz, M., Azizi, L., Buiatti, M., … Dehaene, S. (2015). Disruption of hierarchical predictive coding during sleep. Proceedings of the National Academy of Sciences, 112(11), E1353–E1362.

Uchida, S., Maloney, T., & Feinberg, I. (1992). Beta (20-28 Hz) and delta (0.3-3 Hz) EEGs oscillate reciprocally across NREM and REM sleep. Sleep, 15(4), 352–358.

Vallat, R., Lajnef, T., Eichenlaub, J.-B., Berthomier, C., Jerbi, K., Morlet, D., & Ruby, P. M. (2017). Increased Evoked Potentials to Arousing Auditory Stimuli during Sleep: Implication for the Understanding of Dream Recall. Front Hum Neurosci, 11(132). doi:10.3389/fnhum.2017.00132

van Schalkwijk, F. J., Sauter, C., Hoedlmoser, K., Heib, D. P., Klösch, G., Moser, D., … Schabus, M. (2017). The effect of daytime napping and full-night sleep on the consolidation of declarative and procedural information. Journal of Sleep Research.

Wehrle, R., Kaufmann, C., Wetter, T. C., Holsboer, F., Auer, D. P., Pollmächer, T., & Czisch, M. (2007). Functional microstates within human REM sleep: first evidence from fMRI of a thalamocortical network specific for phasic REM periods. European Journal of Neuroscience, 25(3), 863–871.

Wilcox, R. R. (2011). Introduction to robust estimation and hypothesis testing: Academic press.

Wilcox, R. R., & Tian, T. S. (2011). Measuring effect size: a robust heteroscedastic approach for two or more groups. Journal of Applied Statistics, 38(7), 1359–1368. doi:10.1080/02664763.2010.498507

World Medical Association (WMA). (1964). Ethical Principles for Medical Research Involving Human Subjects. 10/13. Retrieved from http://www.wma.net/en/30publications/10policies/b3/

Wyatt, J. K., Cecco, A. R.-D., Czeisler, C. A., & Dijk, D.-J. (1999). Circadian temperature and melatonin rhythms, sleep, and neurobehavioral function in humans living on a 20-h day. American Journal of Physiology - Regulatory, Integrative and Comparative Physiology, 277(4), R1152–R1163.

